# A convolutional neural network for estimating synaptic connectivity from spike trains

**DOI:** 10.1101/2020.05.05.078089

**Authors:** Daisuke Endo, Ryota Kobayashi, Ramon Bartolo, Bruno B. Averbeck, Yasuko Sugase-Miyamoto, Kazuko Hayashi, Kenji Kawano, Barry J. Richmond, Shigeru Shinomoto

## Abstract

The recent increase in reliable, simultaneous high channel count extracellular recordings is exciting for physiologists and theoreticians because it offers the possibility of reconstructing the underlying neuronal circuits. We recently presented a method of inferring this circuit connectivity from neuronal spike trains by applying the generalized linear model to cross-correlograms. Although the algorithm can do a good job of circuit reconstruction, the parameters need to be carefully tuned for each individual dataset. Here we present another method using a Convolutional Neural Network for Estimating synaptic Connectivity from spike trains (CoNNECT). After adaptation to huge amounts of simulated data, this method robustly captures the specific feature of monosynaptic impact in a noisy cross-correlogram. There are no user-adjustable parameters. With this new method, we have constructed diagrams of neuronal circuits recorded in several cortical areas of monkeys.

## I. INTRODUCTION

More than half a century ago, Perkel, Gerstein, and Moore [1] pointed out that by measuring the influence of one neuron on another through a cross-correlogram, physiologists could infer the strength of the connection between the neurons. If this were done for lots of pairs of neurons, a map of the neuronal circuitry could be built. Now, with the advent of high-quality simultaneous recording from large arrays of neurons, it might have become possible to map the structures of neuronal circuits.

The original cross-correlation method can give plausible inferences about connections. However, in many cases, it also tended to suggest the presence of connections that are spurious, i.e., false positives (FPs). There were many possible sources for the lack of reliability and specificity, such as large fluctuations produced by external signals or higher-order interactions among neurons. Over the years, there have been many attempts to minimize the presence of such spurious connections, by shuffling spike trains [2], by jittering spike times [3–6], or by taking fluctuating inputs into account [7–13]. These, in general, helped eliminate the FPs, but they then tended to be conservative, giving rise to false negatives (FNs), i.e., missing existing connections.

Recently, we developed an estimation method by applying the generalized linear model (GLM) to each cross-correlogram [14]. The estimation method we call GLMCC works well in balancing the conflicting demands of reducing FPs and reducing FNs, demonstrating that the cross-correlogram image actually contains sufficient information from which to infer the presence of monosynaptic connectivity. GLMCC nonetheless has a shortcoming: the estimation results are sensitive to the model parameters, and therefore the parameters need to be tuned for the spiking data.

Here, we develop another method: Convolutional Neural Network for Estimating synaptic Connectivity from spike Trains (CoNNECT). The premise is that a convolutional neural network is good at capturing the features important for distinguishing among different categories of images [15–18]; we apply it to a cross-correlogram, expecting that it is capable of detecting the signature of the monosynaptic impact in the one-dimensional cross-correlogram image. Our new method CoNNECT is easy to use, and it works robustly with data arising from different cortical regions in non-human primates. The convolutional neural network has tens of thousands of internal parameters. The parameters can be adjusted using hundreds of thousands of pairs of spike trains generated with a large-scale simulation of the circuitry of realistic model neurons. To reproduce large fluctuations in real spike trains, we added external fluctuations to the model neurons in the simulation.

CoNNECT promptly provides reasonable inference. It does not, however, give a rationale for why the result was derived, whereas our previous algorithm GLMCC does because it fits an interaction kernel to the cross-correlogram. These methods, therefore, have different strengths and weaknesses and can be used in combination in a complementary manner. Namely, the inference given by CoNNECT can be used for guiding GLMCC to search for suitable parameters, and GLMCC can provide interpretation.

We evaluated the accuracy of estimation by comparing the inference with the true connections, using synthetic data generated by simulating circuitries of model neurons, and compared the performance of CoNNECT with that of GLMCC, as well as the classical cross-correlogram method [19, 20], the Jittering method [4, 5], and an extended GLM method [13]. After confirming the performance of the model, we applied CoNNECT to parallel spike signals recorded from three cortical areas of monkeys and obtained estimation of the local neuronal circuitry among many neurons. We have found that the connections among recorded units are sparse; they are less than 1% for all three datasets.

## II.RESULTS

### A. Training and validating with synthetic data

CoNNECT infers the presence or absence of monosynaptic connections between a pair of neurons and estimates the amplitude of the postsynaptic potential (PSP) that one neuron would drive in another. The estimation is performed by applying a convolutional neural network [15–18] to a cross-correlogram obtained for every pair of spike trains (Fig. 1a). The network has an output layer consisting of two units. One unit indicates the presence or absence of connectivity with a real value *z* ∈ [0, 1] by thresholding at 0.5. Another is the level of PSP represented in a unit of [mV]. The network was trained with spike trains generated by a numerical simulation of a network of multiple-timescale adaptive threshold (MAT) model neurons [21–25] interacting through fixed synapses. In a large-scale simulation, we applied fluctuating inputs to a subset of neurons to reproduce large fluctuations in real spike trains *in vivo* (Fig. 1b). Figures 1 c, d, and e demonstrate sample spike trains, histograms of the firing rates of excitatory and inhibitory neurons [26], and firing irregularity measured in terms of the local variation of the interspike intervals *Lv* [27, 28]. The training data does not contain many low firing rate neurons, considering the situation that low firing units are often discarded when analyzing real data. The details of the learning procedure are summarized in METHODS.

**FIG. 1.**
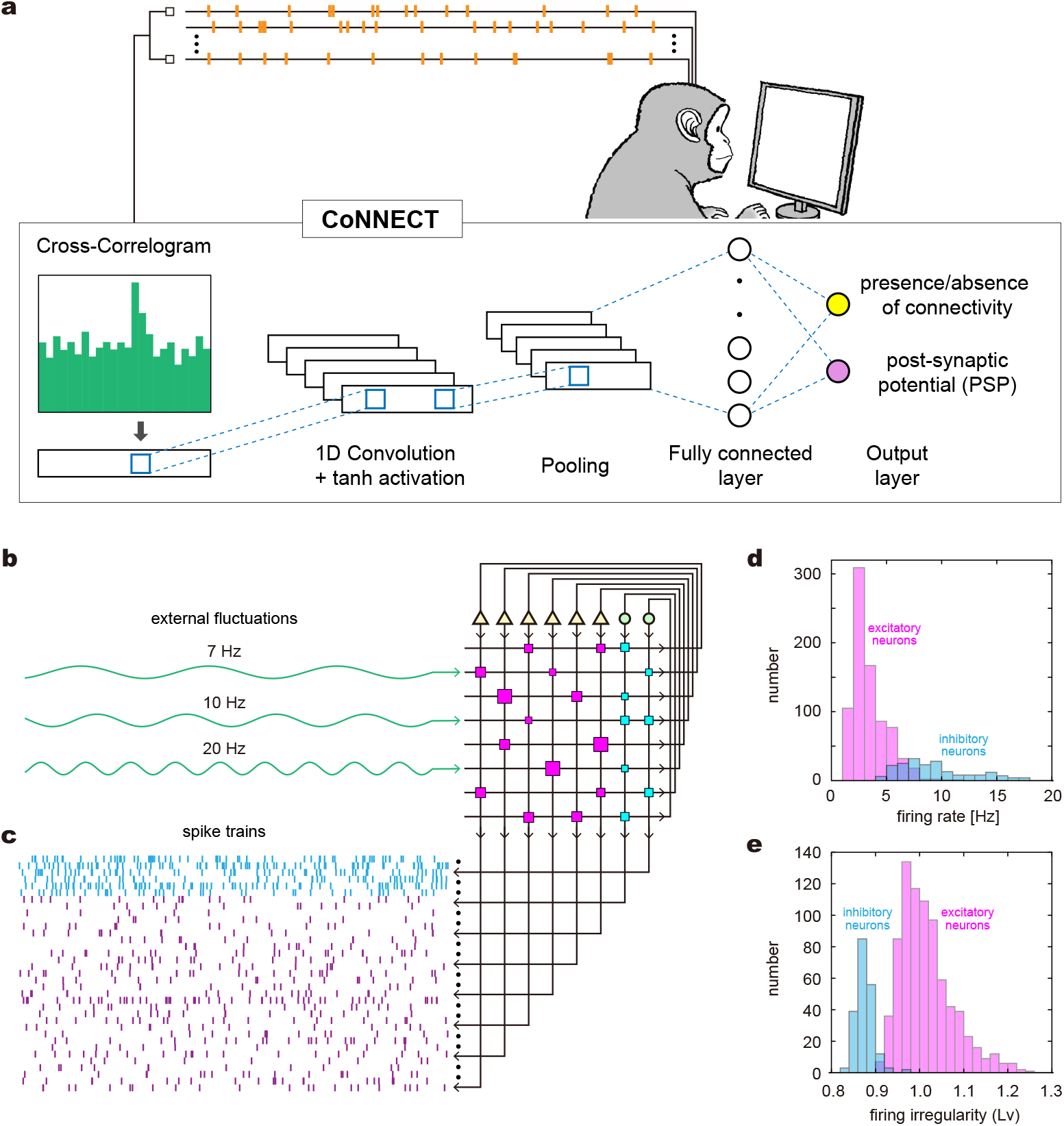
The architecture of CoNNECT. (a) The algorithm infers the presence or absence of monosynaptic connectivity and the value of postsynaptic potential (PSP) from the cross-correlogram obtained from a pair of spike trains. The figure of a monkey was illustrated by Kai Shinomoto and licenced to Springer Nature Limited. (b) The algorithm is trained with spike trains generated by a numerical simulation of neurons interacting through fixed synapses. Slow fluctuations were added to a subset of neurons to reproduce large fluctuations in real spike trains *in vivo*. (c) Sample spike trains (cyan: inhibitory neurons; magenta: excitatory neurons). (d) Firing rates of excitatory and inhibitory neurons. (e) Firing irregularity measured in terms of the local variation of the interspike intervals *Lv*.

We validated the estimation performance of CoNNECT using novel spike trains generated by another neuronal circuit with different connections. Figure 2a depicts an estimated connection matrix, referenced to the true connection matrix, of 50 neurons. Here, the estimation was done with spike trains recorded for 120 min. Of 50 spike trains, 40 and 10 are, respectively, sampled from 800 excitatory and 200 inhibitory neurons. Figure 2b compares the estimated PSPs against true values. We have presented an estimated PSP as being 0 if the connection is not detected. Points lying on the nonzero *x*-axis are existing connections that were not detected or FNs. Points lying on the nonzero *y*-axis are spurious connections assigned for unconnected pairs or FPs. Figure 2c depicts how the numbers of FNs and FPs for excitatory and inhibitory categories changed with the recording duration or the length of spike trains (10, 30, and 120 min). While the number of FPs or spurious connections does not depend largely on the recording duration, the number of FNs or missing connections decreased with the period, implying that more synaptic connections of weaker strength are revealed by increasing the recording time.

**FIG. 2.**
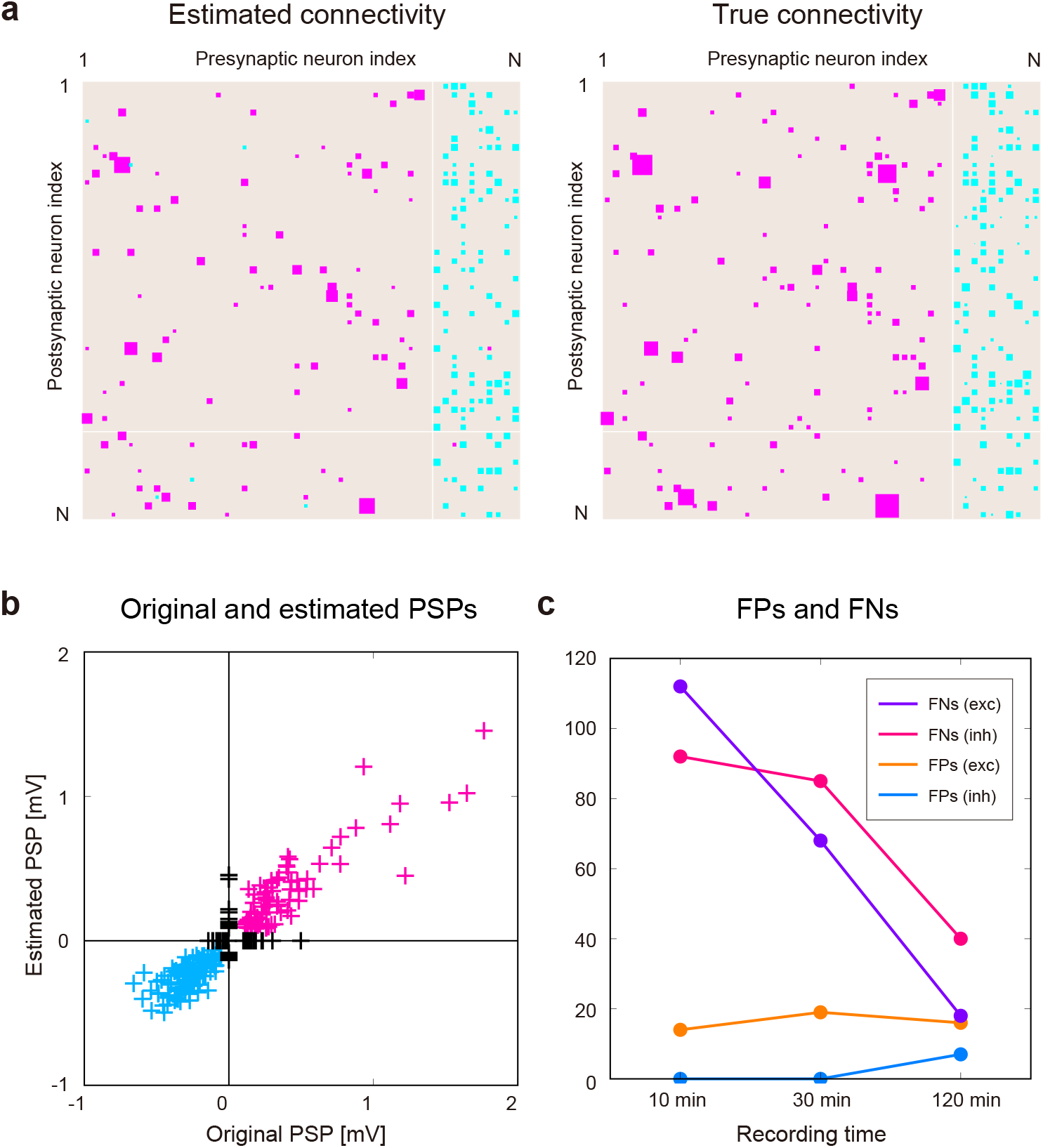
Synaptic connections estimated using CoNNECT. (a) An estimated connection matrix, referenced to a true connection matrix. Of 50 neurons, 40 and 10 are excitatory and inhibitory neurons sampled from 1,000 model neurons simulated for 120 min. Excitatory and inhibitory connections are represented by magenta and cyan squares of the sizes proportional to the postsynaptic potential (PSP). (b) Estimated PSPs plotted against true parameters. Points on the nonzero *y*-axis represent the false positives (FPs) for unconnected pairs. Points on the nonzero *x*-axis represent the false negatives (FNs). (c) The numbers of FPs and FNs for excitatory and inhibitory categories counted for different recording durations.

#### Comparison with other estimation methods

There are many algorithms that were developed to estimate synaptic connections from spike trains [1–13, 19, 29–31]. We compared CoNNECT with the conventional cross-correlation method (CC) [19], Jittering method [4], Extended GLM [13], and GLMCC [21] for their ability to estimate connectivity using synthetic data. Figure 3 shows connection matrices determined by the four methods referenced to the true connection matrices. In the lower panels, we demonstrated the performances in terms of the false positive (discovery) rate (FPR) and false negative (omission) rate (FNR) for excitatory and inhibitory categories; smaller values are better. Here we estimated the mean and SD of the performance by applying each method to 8 test datasets of 50 neurons. Overall performance with FPR and FNR was measured in terms of the Matthews correlation coefficient (MCC) (see METHODS). The MCCs for these estimation methods are shown in the right edge panel; a larger MCC is better. For evaluating the performances, we adopted spiking data generated by a network of MAT models and a network of Hodgkin-Huxley (HH) type models (METHODS). In computing the numbers of FPs and FNs, we ignored small excitatory connections, which are inherently difficult to discern with this observation duration. We took the lower thresholds as 0.1 mV for the MAT simulation and 1 mV for the HH simulation so that the visible connectivity is about 10 %, but the relative performances between different models are unchanged even if we change the thresholds.

**FIG. 3.**
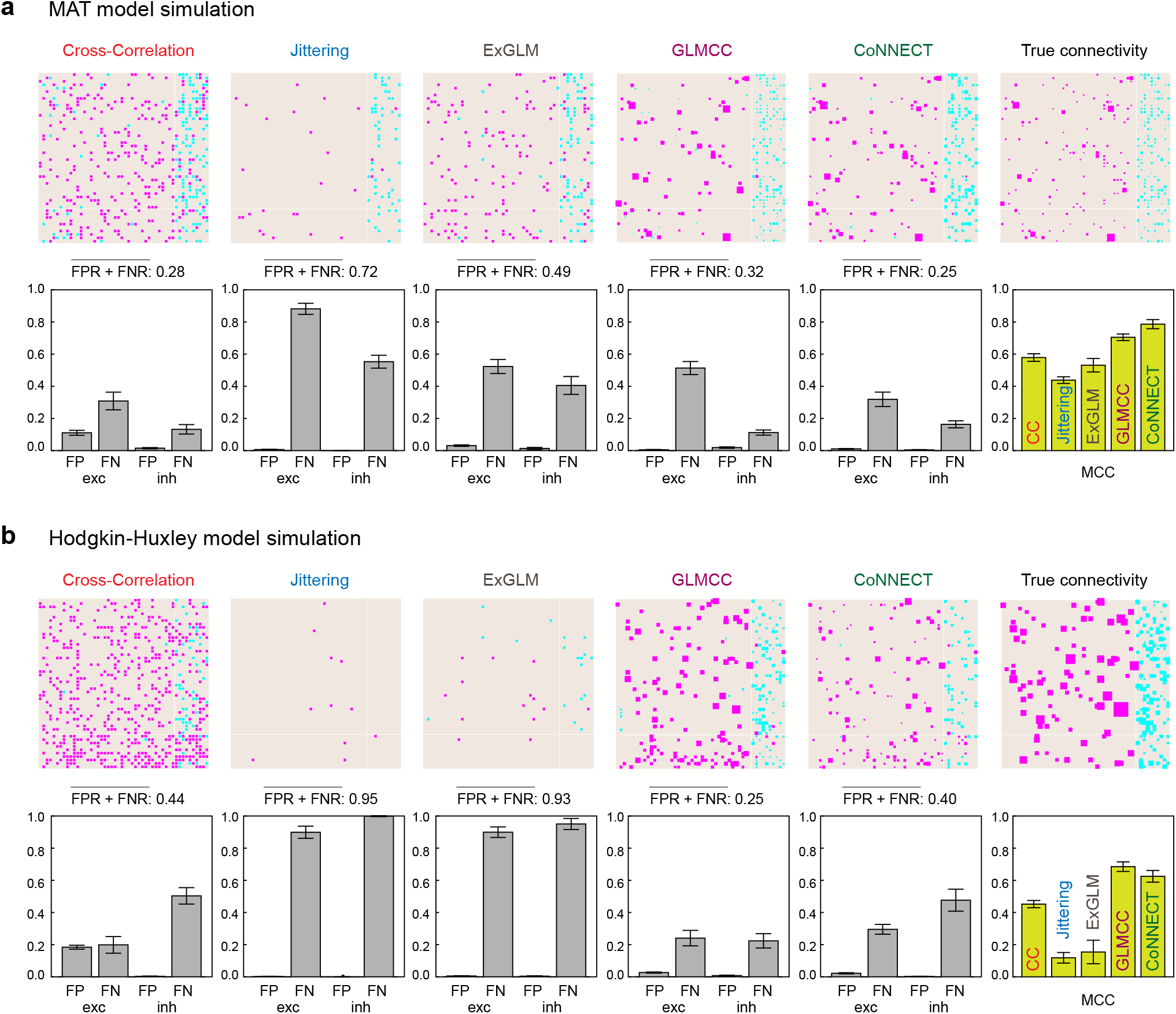
Comparison of estimation methods using two kinds of synthetic data. (a) The multiple-timescale adaptive threshold (MAT) model simulation. (b) The Hodgkin-Huxley (HH) type model simulation. estimated using the conventional cross-correlation method (CC), Jittering method, Extended GLM (ExGLM), GLMCC, and CoNNECT are depicted, referenced to the true connectivity of the synthetic data. Estimated connections are depicted in equal size for the first two methods because they do not estimate the amplitude of PSP. (lower panels) The false-positive rate (FPR) and false-negative rate (FNR) for excitatory and inhibitory categories; smaller values are better. The mean and SD were obtained by applying each method to 8 test datasets of 50 neurons. The sum of FPR and FNR averaged over excitatory and inhibitory categories is presented above each panel. (lower rightmost panel) Overall performances of the estimation methods compared in terms of the Matthews correlation coefficient (MCC); the larger, the better.

The conventional cross-correlation analysis produced many FPs, revealing a vulnerability to fluctuations in cross-correlograms. The Jittering method succeeded in avoiding FPs but missed many existing connections, thus generating many FNs. The Extended GLM method of given parameters was also rather conservative. In comparison to these methods, GLMCC and CoNNECT have better performance, producing a small number of FPs and FNs and a larger MCC value. Here we have modified GLMCC so that it achieves higher performance than the original algorithm [14] by using the likelihood ratio test to determine the statistical significance (METHODS). When comparing these two algorithms, GLMCC was slightly conservative, producing more FNs, while CoNNECT tended to suggest more connections, producing more FPs.

The converted GLMCC was better than CoNNECT for the HH model data (Fig. 3b), but the converse was true for the MAT model data (Fig. 3a). This might be because CoNNECT was trained using the MAT model data of a similar kind, and GLMCC was constructed by considering the HH model simulation. Although the model performance was examined with independent datasets, the HH model simulations would be more similar than across the HH model and MAT model. As the GLMCC parameters were selected with the HH model simulation, it naturally works better for the HH data than MAT simulation data and vice versa.

#### Comparison of different learning conditions

While the convolutional neural network has the advantage that tens of thousands of parameters can be suitably adjusted to reproduce given datasets, it does not guarantee the generalization capability. We evaluated the generalization capability of our convolutional network by changing the number of out-channels representing the degree of system adaptability from 1 to 10. Figure 4a depicts the numbers of FPs and FNs in the above, and the overall performances measured in terms of MCC in the below, which were obtained for the MAT model simulation data (left panel) and the Hodgkin-Huxley model simulation data (right panel). We observe that the convolutional network consisting of 1 channel slightly “under-fits” because of the little flexibility, whereas that of 10 channels slightly over-fits the data, exhibiting slightly lower MCC. Thus we have employed the network of 5 channels, consisting of about fifty thousand parameters.

**FIG. 4.**
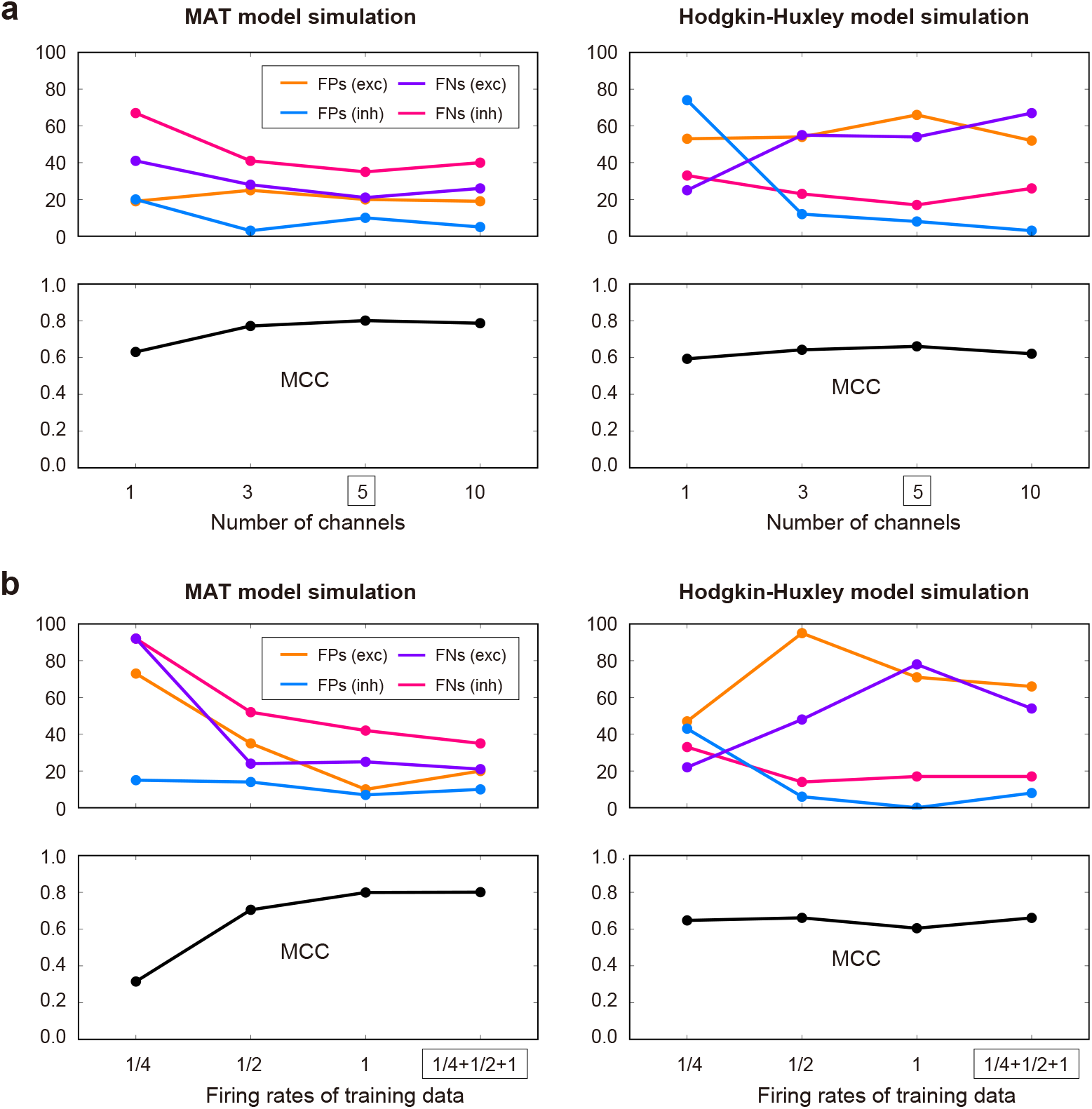
Comparison of different learning conditions. (a) The convolutional networks of different numbers of channels. We have adopted a network of 5 channels. (b) The convolutional networks trained using the cross-correlations rescaled by 1/4 and 1/2 times the original (indicated as “1/4” and “1/2”), the original cross-correlations (“1”), and all the data (“1/4+1/2+1”). We have adopted the network trained with all the data. The numbers of FPs and FNs for the excitatory and inhibitory categories estimating the connectivity of 50 neurons are depicted above, and the Matthews correlation coefficient (MCC) is depicted below. The performances are tested with the MAT model simulation data (left panel) and the Hodgkin-Huxley model simulation data (right panel).

#### Selection of training data sets

To make the convolutional network applicable to data of a wider variety, we have trained the network using the cross-correlograms augmented by rescaling the time. Figure 4b depicts the estimation performances of networks trained using the cross-correlations rescaled by 1/4 and 1/2 times of the original (indicated as “1/4” and “1/2”), the original cross-correlations (“1”), and all the data (“1/4+1/2+1”). The networks trained with lower firing rates exhibited lower performances. We have adopted the network trained with all the data (“1/4+1/2+1”) because it gave the highest performance in estimating connectivity.

#### Cross-correlograms

To observe the situations in which different estimation methods succeeded or failed in detecting the presence or absence of synaptic connectivity, we examined sample cross-correlograms of neuron pairs of a network of MAT model neurons. Figure 5 depicts neuron pairs that exhibited various patterns including pathological cases. The majority of neuron pairs are of successful cases demonstrated in the upper part of the figure. Some cross-correlograms from this simulation exhibited large fluctuations that resemble what is seen in real biological data. These were produced by external fluctuations added to a subset of neurons, making the connectivity inference difficult. The inference results obtained by the four estimation methods are distinguished with colors; magenta, cyan, and gray represent that estimated connections were excitatory, inhibitory, or unconnected, respectively. We also superimposed a GLM function fitted to each cross-correlogram.

**FIG. 5.**
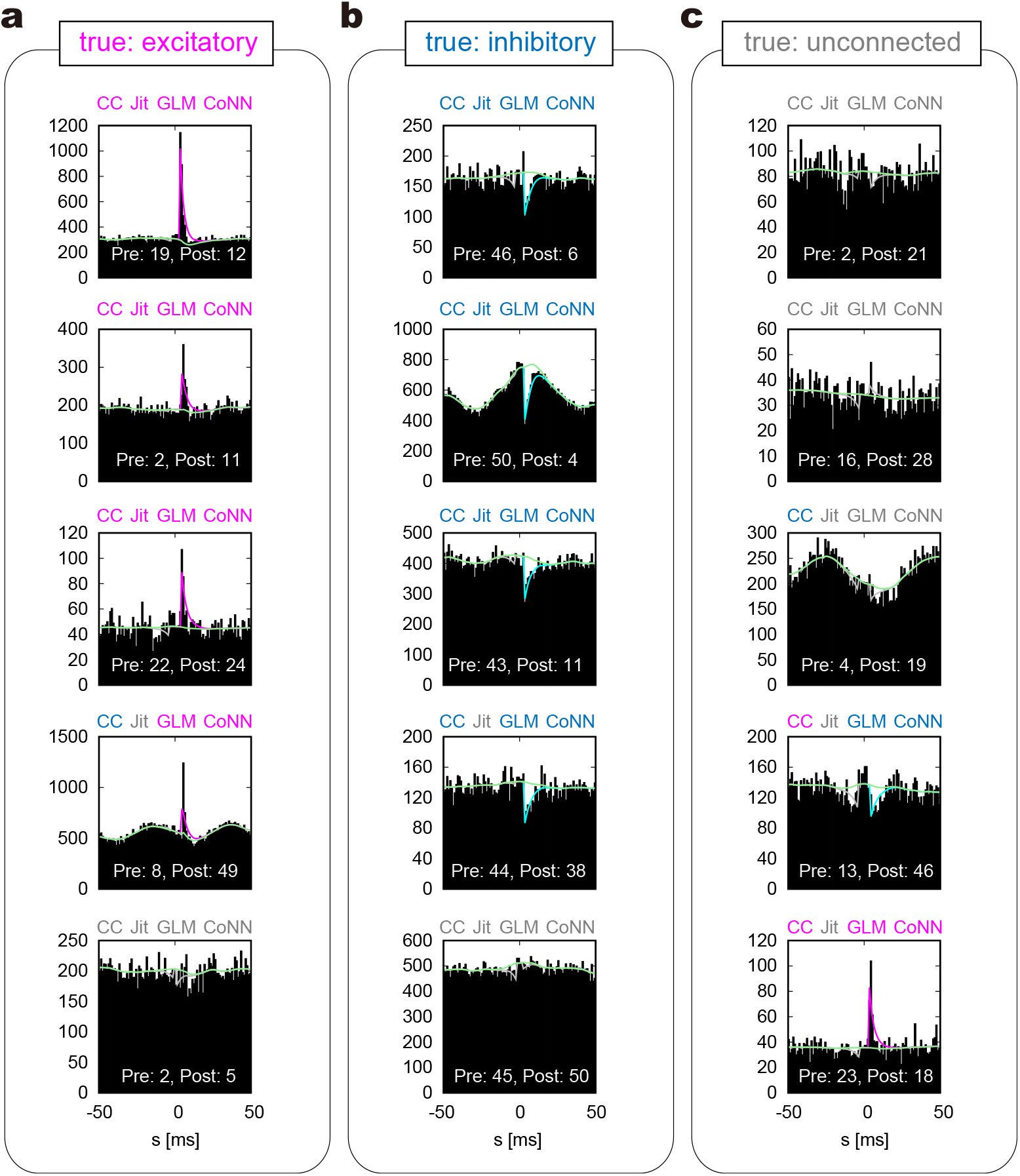
Sample cross-correlograms obtained from the MAT model simulation. (a), (b), and (c) Pairs of neurons that have excitatory and inhibitory connections, and are unconnected, respectively. Four kinds of estimation methods, the cross-correlation (CC), the Jittering (Jit), GLMCC (GLM), and CoNNECT (CoNN), were applied to cross-correlograms. Their estimation (excitatory, inhibitory, and unconnected) are respectively distinguished with colors (magenta, cyan, and gray). The lines plotted on the cross-correlograms are the GLM functions fitted by GLMCC. The causal impact from a pre-neuron to a post-neuron appears on the right half in each cross-correlogram.

Figures 5a and 5b depict sample cross-correlograms of neuron pairs that are connected with excitatory and inhibitory synapses, respectively. For the first three cross-correlograms from the top, all four estimation methods succeeded in detecting excitatory or inhibitory connections, thus making true positive (TP) estimations. For the fourth case, the Jittering method failed to detect the connection. This implies that the Jittering method is rather conservative for producing FPs, and as a result, has produced many FNs. In this case, the cross-correlation method (CC) has mistaken the excitatory synapse as inhibition due to the large wavy fluctuation in the cross-correlogram. For the last cases, all four estimation methods failed to detect the connection, resulting in FNs. This would have been because the original connections were not strong enough to produce significant impacts on the cross-correlograms.

Figure 5c depicts sample cross-correlograms of unconnected pairs. For the first two crosscorrelograms, all four estimation methods judge the absence of connections correctly (or the null hypothesis of the absence of connection was not rejected), resulting in true negatives (TNs). For the third pair, the CC suggested the presence of a connection, resulting in an FP. This demonstrates that the conventional cross-correlation method is fragile in the presence of large fluctuations. For the fourth and the last cases, the CC, GLMCC, and CoNNECT have suggested monosynaptic connections. The sharp peaks appearing in the cross-correlogram would have been caused by indirect interaction via other neurons. In such cases, however, it is difficult to discern the absence of a monosynaptic connection solely from the cross-correlogram.

### B. Analyzing experimental data

We examined spike trains recorded from the prefrontal (PF), inferior temporal (IT), and the primary visual (V1) cortices of monkeys using the Utah arrays. Experimental conditions of individual data are summarized in METHODS. Because neurons with low firing rates do not provide enough evidence for the connectivity, we have excluded low firing units and examined those that have fired more than 1 Hz.

#### Preprocessing experimental data

Some of the experimentally available cross-correlograms exhibit a sharp drop near the origin for a few ms due to the shadowing effect, in which near-synchronous spikes cannot be detected [32]. This effect disrupts the estimation of synaptic impacts that should appear near the origin of the cross-correlogram. The data were obtained with a sorting algorithm specifically used for the Utah array exhibit rather broad shadowing effects larger than 1 ms (up to 1.75 ms). Here, we analyzed the experimental data by removing an interval of 0 *±* 2 ms in the cross-correlogram and applying the estimation method to a cross-correlogram obtained by concatenating the remaining left and right parts (Figs. 6a and b). We also conducted this operation in the analysis of synthetic data.

**FIG. 6.**
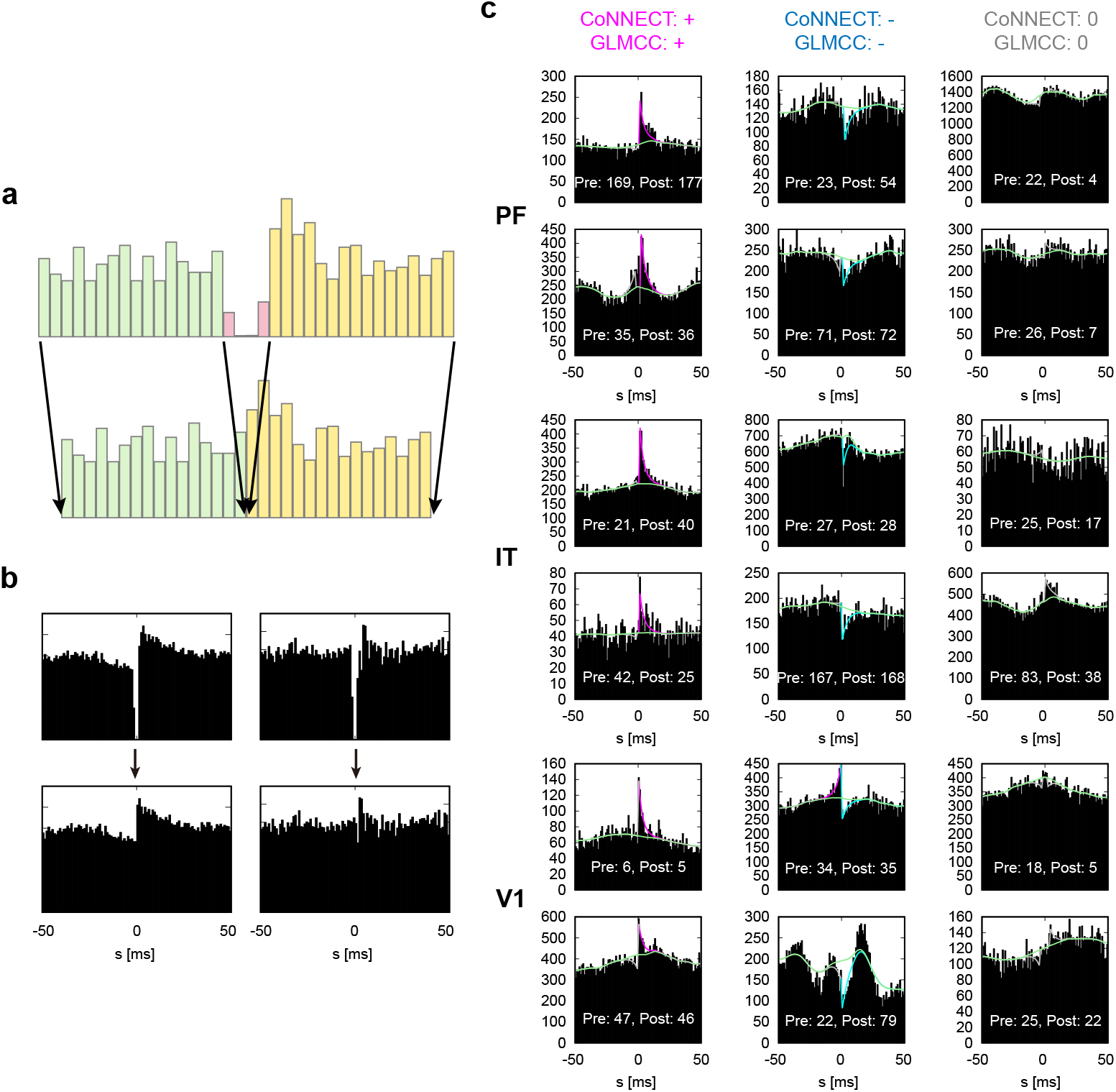
Cross-correlograms of real spike trains recorded from PF, IT, and V1 using the Utah arrays. (a) An interval of 0 *±* 2 ms in the original cross-correlogram was removed to mitigate the shadowing effect, in which near-synchronous spikes were not detected. (b) Processing real cross-correlograms. (c) The cross-correlograms for which CoNNECT and GLMCC gave the same inference. The fitted GLM functions are superimposed on the histograms. The causal impact from a pre-neuron to a post-neuron appears on the right half in the cross-correlogram.

Figure 6c demonstrates the cross-correlograms of sample neuron pairs for which both CoNNECT and GLMCC estimated connections which were excitatory, inhibitory, or absent (unconnected). It was observed that the real cross-correlograms are accompanied by large fluctuations. Nevertheless, CoNNECT and GLMCC are able to detect the likely presence or absence of synaptic interaction by ignoring the severe fluctuations.

#### Connection matrices

Figure 7 depicts the estimated connections for the entire three datasets of PF, IT, and V1. The units in the connection matrices are arranged in the order provided by a sorting algorithm, and accordingly, units of neighboring indexes of the matrices tended to have been spatially closely located. All three connection matrices had more components in near diagonal elements, implying that neurons in a nearby location are more likely to be connected. The firing rate and irregularity (the local variation of the interspike intervals *Lv* [27, 28]) are shown in the rightmost panels. The summary statistics in Table I reflect differences in firing rate between excitatory and inhibitory cells in PF and IT but not V1. The firing irregularity of excitatory neurons is slightly higher than that of inhibitory neurons, consistent with the previous results.

**TABLE I.**
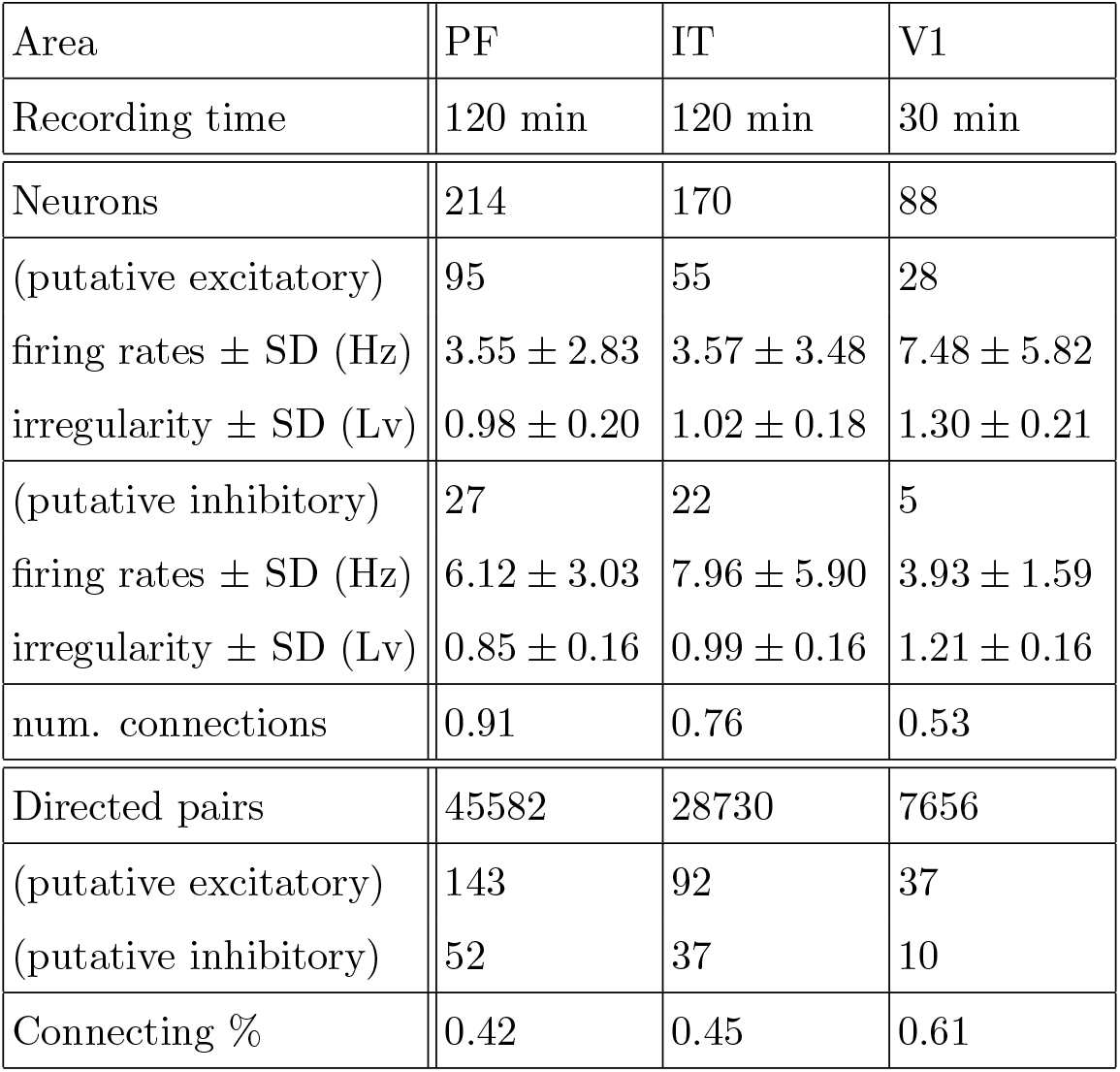
Results of analyzing experimental datasets.

| Area | PF | IT | V1 |
| --- | --- | --- | --- |
| Recording time | 120 min | 120 min | 30 min |
| Neurons | 214 | 170 | 88 |
| (putative excitatory) | 95 | 55 | 28 |
| firing rates ± SD (Hz) | $3.55 \pm 2.83$ | $3.57 \pm 3.48$ | $7.48 \pm 5.82$ |
| irregularity ± SD (Lv) | $0.98 \pm 0.20$ | $1.02 \pm 0.18$ | $1.30 \pm 0.21$ |
| (putative inhibitory) | 27 | 22 | 5 |
| firing rates ± SD (Hz) | $6.12 \pm 3.03$ | $7.96 \pm 5.90$ | $3.93 \pm 1.59$ |
| irregularity ± SD (Lv) | $0.85 \pm 0.16$ | $0.99 \pm 0.16$ | $1.21 \pm 0.16$ |
| num. connections | 0.91 | 0.76 | 0.53 |
| Directed pairs | 45582 | 28730 | 7656 |
| (putative excitatory) | 143 | 92 | 37 |
| (putative inhibitory) | 52 | 37 | 10 |
| Connecting % | 0.42 | 0.45 | 0.61 |

**FIG. 7.**
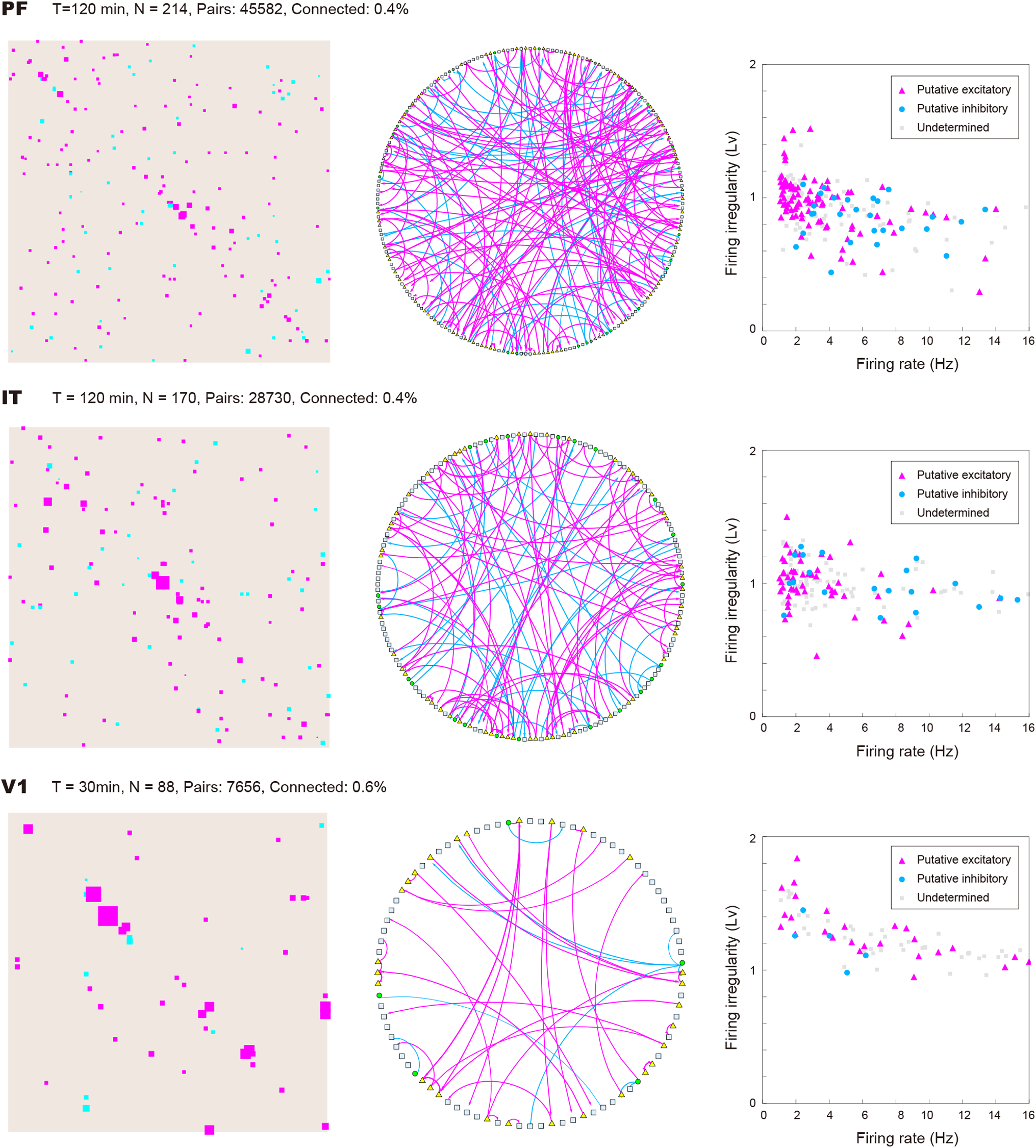
Connection matrices and diagrams estimated for spike trains recorded from the prefrontal (PF), inferior temporal (IT), and the primary visual (V1) cortices of monkeys. In the connection diagrams, excitatory and inhibitory dominant units are depicted as triangles and circles, respectively, and units with no outgoing connections or those that innervate equal numbers of estimated excitatory and inhibitory connections are depicted as squares. (rightmost panels) The firing rate and irregularity (*L*_*v*_) of the putative excitatory and inhibitory units identified by the E-I dominance index. Units that had no innervating connections or those that exhibited vanishing E-I dominance indices are depicted in gray.

Table I summarizes the statistics of the three datasets. Each neuron is assigned as putative excitatory, putative inhibitory, or undetermined, according to whether the excitatory–inhibitory (E–I) dominance index is positive, negative, or undetermined (or zero), respectively. Here, the E–I dominance index is defined as *d*_ei_ = (*n*_e_ − *n*_i_)*/*(*n*_e_ + *n*_i_), in which *n*_e_ and *n*_i_ represent the numbers of excitatory and inhibitory identified connections projecting from each unit, respectively [14]. The row “num. connections” indicates the average number of innervated connections per neuron.

Because the number of innervated connections for each neuron is only a few, the majority of *d*_*ei*_ is either 0 or 1. Though we have obtained many connections, the total number of all pairs is enormous, scaling with the square of the number of units, and accordingly, the connectivity is sparse (less than 1% for each (directed) pair of neurons).

In contrast to synthetic data, the currently available experimental data do not contain information regarding the true connectivity. To examine the stability of the estimation, we split the recordings in half and compared estimated connections from each half. If the real connectivity is stable, we may expect the estimated connections have overlap between the first and second halves. Figure 8a represents the connection matrices obtained from the first and second halves of the spike trains recorded from PF, IT, and V1. Figure 8b compares the estimated PSPs in two periods. Many estimated connections appear only on one of the two. This might be simply due to statistical fluctuation or due to real changes in synaptic connectivity. Nevertheless, it may be noteworthy that the excitatory connections of large amplitudes were detected relatively consistently between the first and second halves. Namely, they appear in the first and third quadrants diagonally, implying that they have the same signs with similar amplitudes.

**FIG. 8.**
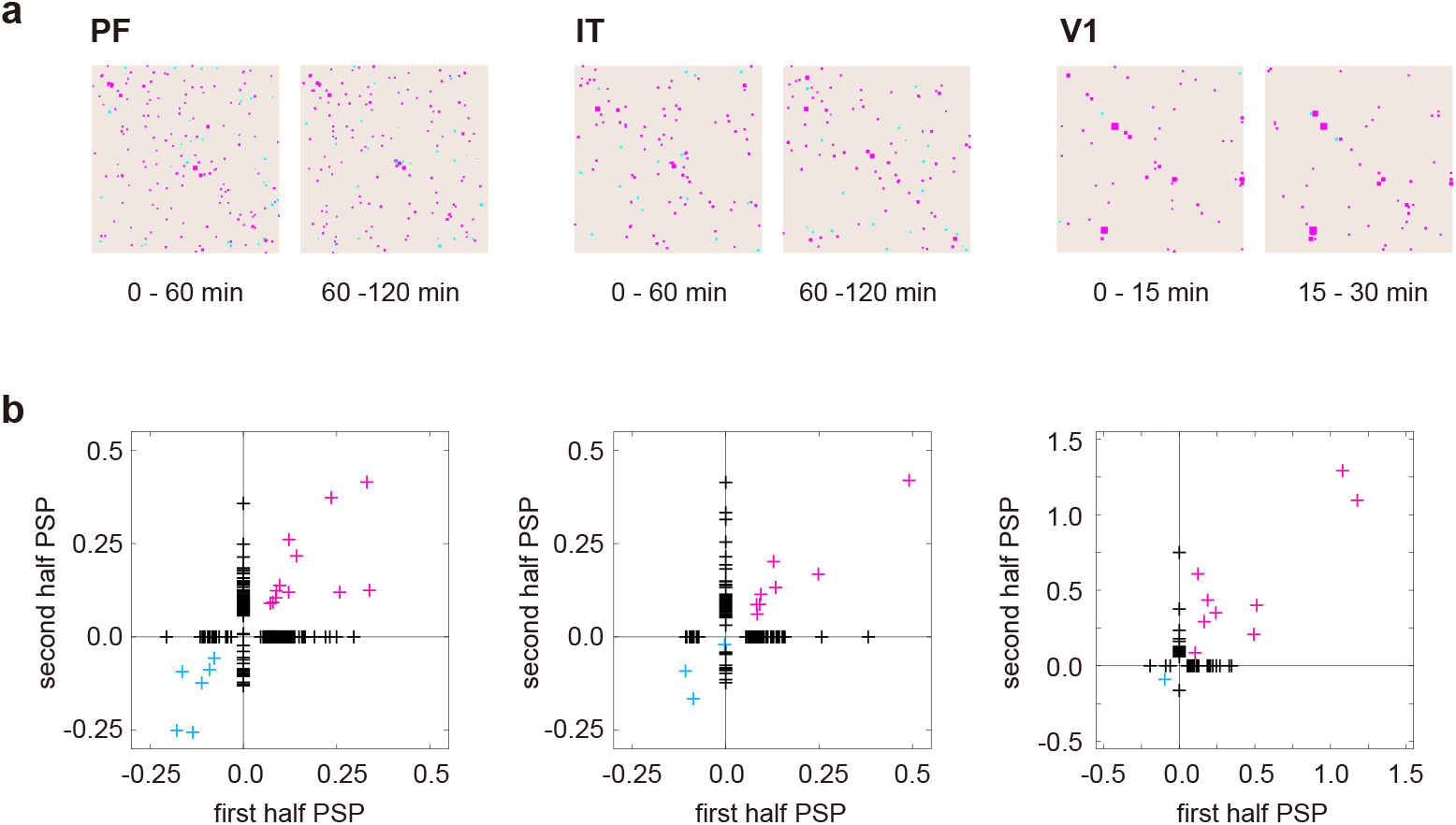
Stability of connection estimation. (a) Connection matrices estimated for the first and second halves of the spike trains recorded from PF, IT, and V1. (b) Comparison of the PSPs estimated from the first and second halves.

## III. DISCUSSION

Here we have devised a new method for estimating synaptic connections based on a convolutional neural network. While this method does not require adjusting parameters for individual data, it robustly provides a reasonable estimate of synaptic connections by tolerating large fluctuations in the data. This high performance was obtained by training a convolutional neural network using a considerable amount of training data generated by simulating a network of model spiking neurons subject to fluctuating current.

We compared CoNNECT with the conventional cross-correlation method, the Jittering method, Extended GLM, and GLMCC in their ability to estimate connectivity, using synthetic data obtained by simulating neuronal circuitries of fixed synaptic connections. Both CoNNECT and GLMCC exhibited high performance in predicting individual synaptic connections, superior to other methods.

Then we applied CoNNECT to simultaneously recorded spike trains recorded from monkeys using the Utah arrays. We have found that the connections among recorded units are sparse; they are less than 1% for all three datasets. To test the reliability of the estimation, we divided the entire recording interval in half and estimated connections for respective intervals. We have seen that strong excitatory connections overlap between the periods. This result implies that the estimation is reliable for the strong connectivity, and the connectivity lasts at least for hours.

The cross-correlograms of real biological data (Fig. 6) turned out to be even more complicated than those of synthetic data (Fig. 5), which were generated by adding large fluctuations to individual neurons (Fig. 1). The complicated features in real cross-correlograms were not only due to fluctuations in real circuitry but also due to the sorting algorithm. The most severe bottleneck in estimating connectivity may have been the shadowing effect of a few ms, in which near-synchronous spikes were not detected (Fig. 6a); this effect might hide the first part of a monosynaptic impact, which is expected to show up in a few ms in a cross-correlogram. If the sorting algorithm is improved such that the shadowing duration is shortened, the estimation might be more reliable.

In this study, we have employed the convolutional neural network to capture the specific signature of monosynaptic impact in a cross-correlogram image. While the convolutional network is known to be robust against the translation of images, the monosynaptic impact is expected to appear at a specific location in the cross-correlogram, particularly exhibiting the delay of a few milliseconds. Thus it might be an interesting challenge to search for other learning algorithms that utilize such information and perform better than the convolutional network.

We used data augmentation technique [33] to increase the number of training examples artificially. Data augmentation is known to improve the performance on various tasks in computer vision [18, 34] and acoustic signal processing [35, 36]. Here we augmented the cross-correlation data by rescaling the time to capture the diverse synaptic interactions. This augmentation also improved the estimation performance of synaptic connectivity (Fig. 4b). Recently, several authors proposed a more systematic approach for data augmentation, e.g., generating augmented data using generative adversarial networks (GANs) [37, 38] and learning the data augmentation policy [39]. Though these approaches focus on the image classification task and require a vast computational resource, it may be interesting to apply these techniques to pursue an advanced data augmentation method for synaptic connectivity estimation.

So far, we have little knowledge about neuronal circuitry in the brain. By collecting more data from high channel count recordings and applying these reliable analysis methods to them, we shall be able to obtain information about neuronal circuitry in different brain regions and learn about network characteristics and the information flow in each area. Ultimately, we expect that we will characterize the network characteristics of different brain regions processing various kinds of information.

## IV. METHODS

### A. Configuration of a neural network for estimating synaptic connectivity

Here we describe the details of a four-layered convolutional neural network [15–18] applied to cross-correlograms obtained for every pair of spike trains to estimate the presence or absence of a connection and its postsynaptic potential (PSP) (Fig. 1). The neural network learns to find a bump or dent in the cross-correlogram caused by a monosynaptic connection.

In particular, the input consists of 100 integer values of the spike counts in a cross-correlation histogram in an interval of [-50, 50] ms with 1 ms bin size. The network comprises a 1-dimensional convolution layer, the average pooling, and the internal layer of fully connected 100 nodes. The output layer consists of two units; one indicates the presence or absence of connectivity with a real value *z* ∈ [0, 1]. Another is the level of PSP represented in a unit of [mV].

#### Training the convolutional neural network

We ran a numerical simulation of a network of 1,000 neurons interacting through fixed synapses in various conditions and trained the neural network with spike trains from 400 units selected from the entire network. Thus, we constructed cross-correlograms of about 80, 000 pairs, each assigned with the teaching signals consisting of the true information about the presence or absence of connectivity (respectively represented as *z* = 1 or 0) and its PSP value in either direction. The training was performed using an algorithm for first-order gradient-based optimization of stochastic objective functions, based on adaptive estimates of lower-order moments, named Adam [40]. The parameters adopted in the learning are summarized in Table II. Details of the architecture are summarized in Table III. Figure 9 demonstrates a set of convolutional kernels that were learned with training data. From the set of learned kernels, we can see some specific features of monosynaptic impact of a few milliseconds appearing in a cross-correlogram. It is also interesting to see a kernel exhibiting a roughly monotonic gradient (the second panel from the left). This might have worked for detrending the large slow fluctuations in the cross correlogram produced by our simulation, which aimed at reproducing real situations.

**TABLE II.**
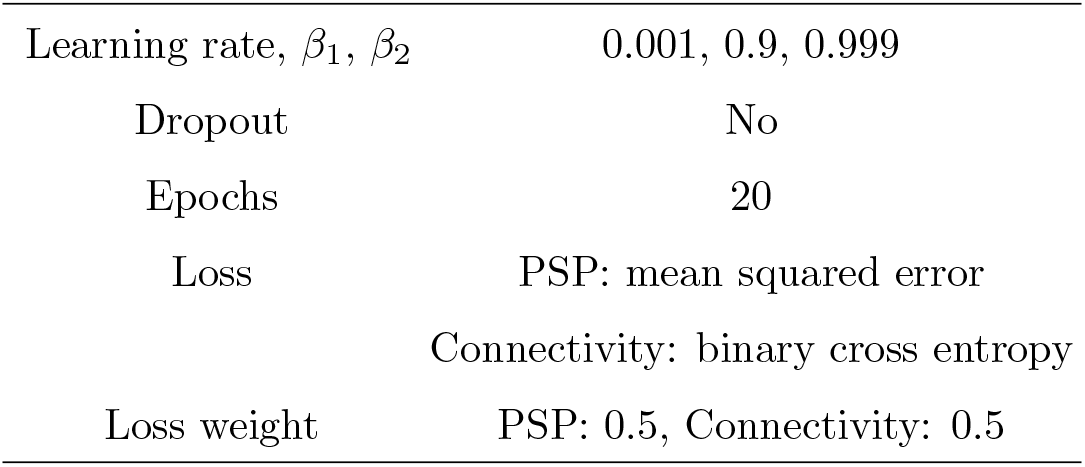
Hyperparameters of the convolutional neural network.

**TABLE III.** Architecture of the convolutional neural network.

| Convolution layer |  |
| --- | --- |
| Kernel size | 10 |
| Number of out-channels | 5 |
| Slide range | 1 |
| Activation function | tanh |
| Hidden layer |  |
| Number of nodes | 100 |
| Activation function | ReLU |

**FIG. 9.**
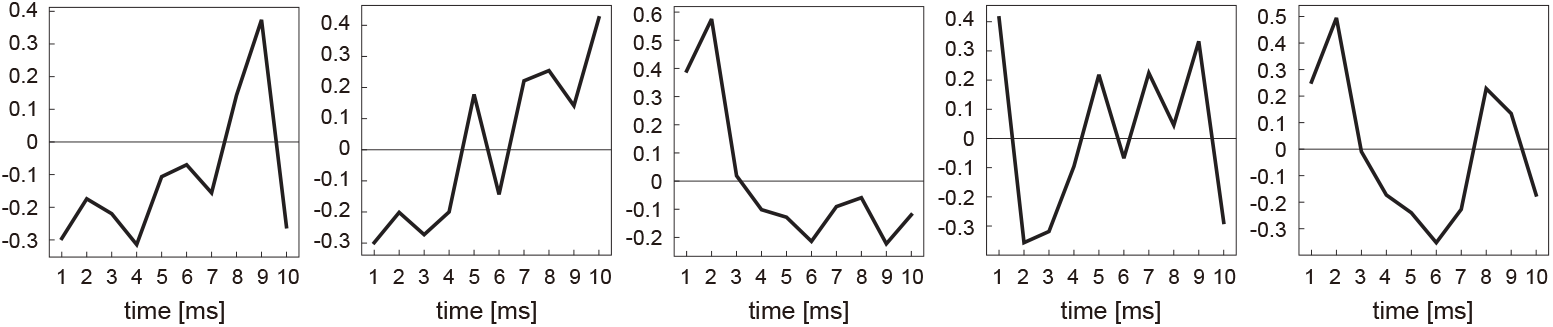
A set of convolutional kernels that were learned from training data. The range of the kernel is 10 ms.

#### Data Augmentation

To make the estimation method applicable to data of a wider range, we performed data augmentation [18, 34–37] (see [33] for review). Namely, we augmented the data by rescaling the cross-correlations by 2 and 4 times and used all the data, including the original data in the learning.

#### Web-application program

A ready-to-use version of the web application, the source code, and example data sets are available at our website, https://s-shinomoto.com/CONNECT/ and are also hosted publicly on Github, accessible via https://github.com/shigerushinomoto. The simulation code is also available at this Github.

### B. Improvement of GLMCC

#### Original framework of GLMCC

In the previous study [14], we developed a method of estimating the connectivity by fitting the generalized linear model to a cross-correlogram, GLMCC. We designed the GLM function as

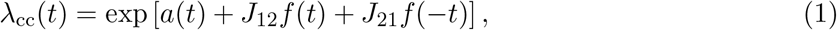

where *t* is the time from the spikes of the reference neuron. *a*(*t*) represents large-scale fluctuations in the cross-correlogram in a window [-*W, W*] (*W* = 50 ms). By discretizing the time in units of Δ(= 1ms), *a*(*t*) is represented as a vector 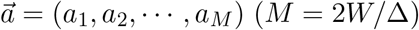. *J*_12_ (*J*_21_) represents a possible synaptic connection from the reference (target) neuron to the target (reference) neuron. The temporal profile of the synaptic interaction is modeled as 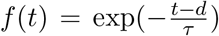 for *t > d* and *f*(*t*) = 0 otherwise, where *τ* is the typical timescale of synaptic impact and *d* is the transmission delay. Here we have chosen *τ* = 4 ms, and let the synaptic delay *d* be selected from 1, 2, 3, and 4 ms for each pair.

The parameters 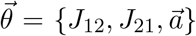 are determined with the maximum *a posteriori* (MAP) estimate, that is, by maximizing the posterior distribution or its logarithm:

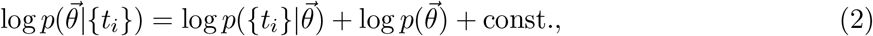

where *{t*_*i*_*}* are the relative spike times. The log-likelihood is obtained as

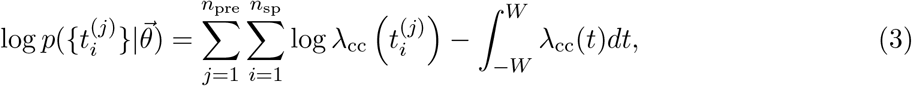

where *n*_pre_ is the number of spikes of presynaptic neuron (*j*). Here we have provided the prior distribution of 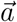 that penalizes a large gradient of *a*(*t*) and uniform prior for *{J*_12_, *J*_21_*}*

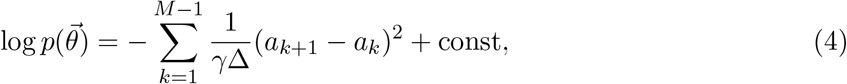

where the hyperparameter *γ* representing the degree of flatness of *a*(*t*) was chosen as *γ* = 2 *×* 10^−4^ [ms^−1^].

#### Likelihood ratio test

The likely presence of the connectivity can be determined by disproving the null hypothesis that a connection is absent. In the original model, this was performed by thresholding the estimated parameters with |*ĵ*_*ij*_ | *>* 1.57*z*_*α*_(*τ T λ*_pre_*λ*_post_)^−1*/*2^, where *z*_*α*_, *T, λ*_pre_, and *λ*_post_ are a threshold for the normal distribution, recording time, firing rates of pre-and post-synaptic neurons. But we realized that this thresholding method might induce a large asymmetry in detectability between excitatory and inhibitory connections.

Instead of a simple thresholding, here we introduce the likelihood ratio test that is a general method for testing hypothesis (Chapter 11 of [41], see also [42]): We compute the likelihood ratio between the presence of the connectivity *J*_*ij*_ = *ĵ*_*ij*_ and the absence of connectivity *J*_*ij*_ = 0 or its logarithm:

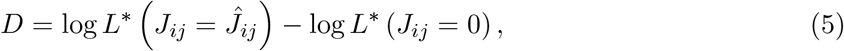

where *L*^*∗*^ (*J*_*ij*_ = *c*) in each case is the likelihood obtained by optimizing all the other parameters with the constraint of *J*_*ij*_ = *c*. It was proven that 2*D* obey the *χ*^2^ distribution in a large sample limit (Wilks’ theorem) [43]. Accordingly, we may reject the null hypothesis if 2*D > z*_*α*_, where *z*_*α*_ is the threshold of *χ*^2^ distribution of a significance level *α*. Here we have adopted *α* = 10^−4^.

### C. Model validation

The performance of CoNNECT was evaluated using the synthetic data generated by independent simulations. The presence or absence of connectivity in each direction is decided by whether or not an output value *z* ∈ [0, 1] exceeds a threshold *θ*. It is possible to reduce the number of FPs by shifting the threshold *θ* to a high level. But this operation may produce many FNs, making many existing connections be missed. To balance the false-positives and false-negatives, we considered maximizing the Matthews correlation coefficient (MCC) [44], as has been done in our previous study [14]. The MCC is defined as

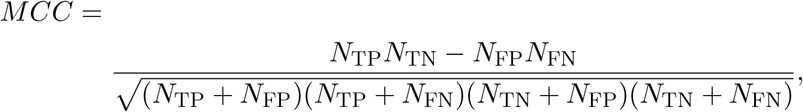

where *N*_TP_, *N*_TN_, *N*_FP_, and *N*_FN_ represent the numbers of true positive, true negative, false positive, and false negative connections, respectively.

We have obtained two coefficients for excitatory and inhibitory categories and taken the macro-average MCC that gives equal importance to these categories (Macro-average) [45], *MCC* = (*MCC*_E_ + *MCC*_I_)*/*2 as we have done in the previous study [14]. In computing the coefficient for the excitatory category *MCC*_E_, we classify connections as excitatory or other (unconnected and inhibitory); for the inhibitory category *MCC*_I_, we classify connections as inhibitory or other (unconnected and excitatory). Here we evaluate *MCC*_E_ by considering only excitatory connections of reasonable strength (EPSP *>* 0.1 mV for the MAT simulation and *>* 1 mV for the HH simulation).

We have confirmed that the Matthews correlation coefficient exhibits a wide peak at about *θ* ∼ 0.5 (Fig. 10), and accordingly, we adopted *θ* = 0.5 as the threshold.

**FIG. 10.**
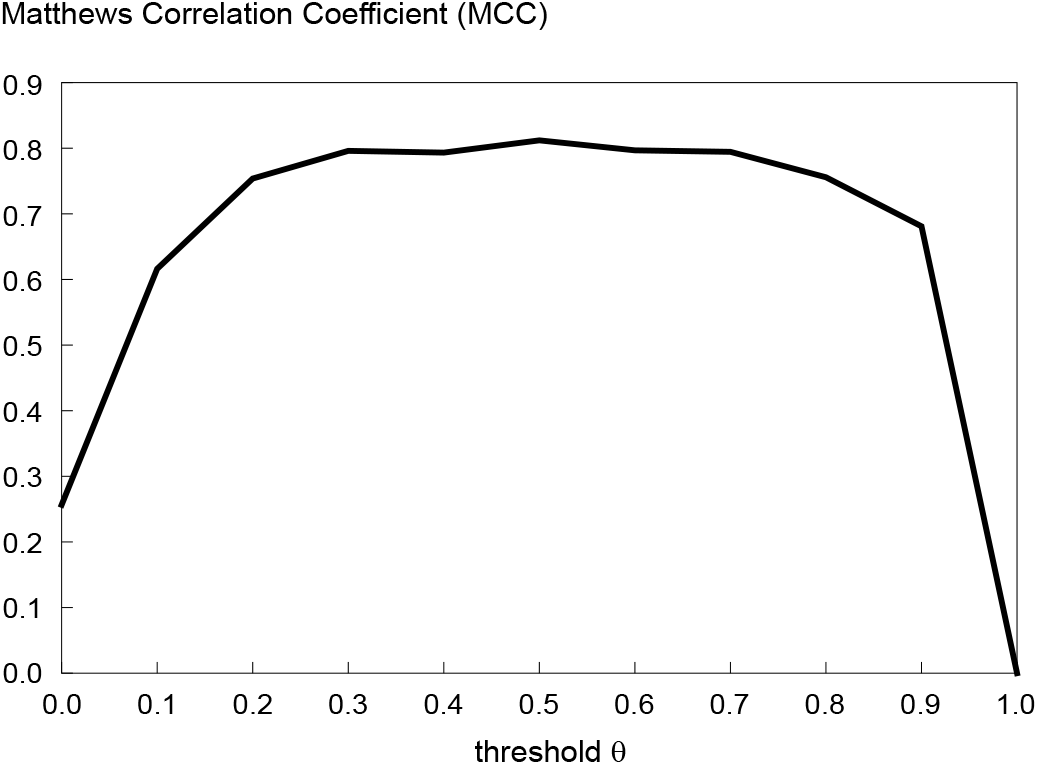
The Matthews correlation coefficient (MCC) plotted against the threshold *θ* for determining the presence and absence of the connection.

### D. A large-scale simulation of a network of MAT neurons

To obtain a large number of spike trains that have resulted under the influence of synaptic connections between neurons, we ran a numerical simulation of a network of 1,000 model neurons interacting through fixed synapses. Of them, 800 excitatory neurons innervate to 12.5 % of other neurons with EPSPs that are log-normally distributed [14, 46–49], whereas 200 inhibitory neurons innervate randomly to 25 % of other neurons with IPSPs that are normally distributed.

#### Neuron model

As for the spiking neuron model, we adopted the MAT model, which is superior to the Hodgkin-Huxley model in reproducing and predicting spike times of real biological neurons in response to fluctuating inputs [21, 23]. In addition, its numerical simulation is stable and fast. The membrane potential of each neuron obeys a simple relaxation equation following the input signal:

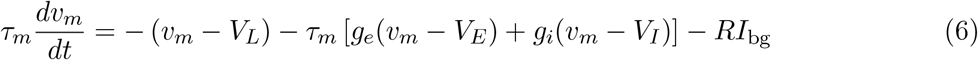

where *g*_*e*_, *g*_*i*_ represents the excitatory conductance and the inhibitory conductance, respectively. Here *RI*_bg_ represent the background noise. The conductance evolves with the

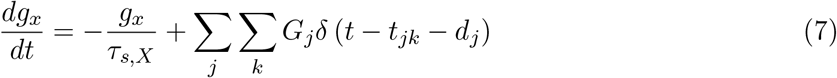

where *τ*_*s*_ is the decay constant, *t*_*jk*_ is the *k*th spike time of *j*th neuron, *d*_*j*_ is a synaptic delay and *G*_*j*_ is the synaptic weight from *j*th neuron. *δ*(*t*) is the Dirac delta function.

Next, the adaptive threshold of each neuron *θ* (*t*) obeys the following equation:

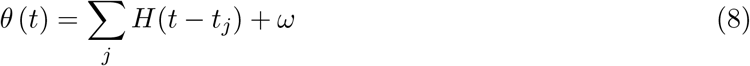

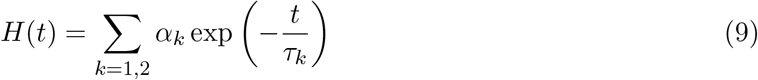

where *t*_*j*_ is the *j*th spike time of a neuron, *ω* is the resting value of the threshold, *τ*_*k*_ is the *k*th time constant, and *α*_*k*_ is the weight of the *k*th component. The parameter values are summarized in Table IV.

**TABLE IV.** Parameters for Neuron Models

| Membrane dynamics |  |
| --- | --- |
| $\tau_m^{\text{excitatory}}, \tau_m^{\text{inhibitory}}$ [ms] | 20, 10 |
| $V_L, V_E, V_I$ [mV] | -70, 0, -80 |
| $\tau_{s,e}, \tau_{s,i}$ [ms] | 1, 2 |
| Threshold dynamics |  |
| $\tau_1, \tau_2$ [ms] | 10, 200 |
| $\omega^{\text{excitatory}}, \omega^{\text{inhibitory}}$ [mV] | -55, -57 |
| $\alpha_1^{\text{excitatory}}, \alpha_2^{\text{excitatory}}$ [mV] | Gauss(1.5, 0.25), 0.5 |
| $\alpha_1^{\text{inhibitory}}, \alpha_2^{\text{inhibitory}}$ [mV] | 3, 0 |

#### Synaptic connections

We ran a simulation of a network consisting of 800 pyramidal neurons and 200 interneurons interconnected with a fixed strength. Each neuron receives 100 excitatory inputs randomly selected from 800 pyramidal neurons and 50 inhibitory inputs selected from 200 interneurons. The excitatory and inhibitory synaptic connections were sampled from respective distributions so that the resulting EPSPs and IPSPs are similar to the distributions adopted in our previous study [14]. In particular, the excitatory conductances 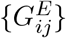 were sampled independently from a log-normal distribution [46, 47].

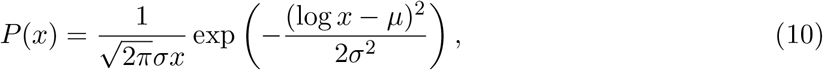

where *µ* = −5.543 and *σ* = 1.30 are the mean and SD of the natural logarithm of the conductances.

The inhibitory conductances 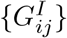 were sampled from the normal distribution:

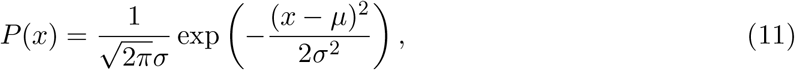

where *µ* = 0.0217 mS cm^−2^, *σ* = 0.00171 mS cm^−2^ are the mean and SD of the conductances. If the sampled value is less than zero, the conductance is resampled from the same distribution. The delays of the synaptic connections from excitatory neurons are drawn from a uniform distribution between 3 and 5 ms. The delays of the synaptic connections from inhibitory neurons are drawn from a uniform distribution between 2 and 4 ms.

#### Background noise

Because our model network is smaller than real mammalian cortical networks, we added a background current to represent inputs from many neurons, as previously done by Destexhe et al.[11, 50].

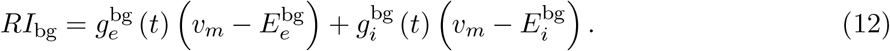

The summed conductance *RI*_bg_ represents random bombardments from a number of excitatory and inhibitory neurons. The dynamics of excitatory or inhibitory conductances can be approximated as a stationary fluctuating process represented as the Ornstein–Uhlenbeck process [51],

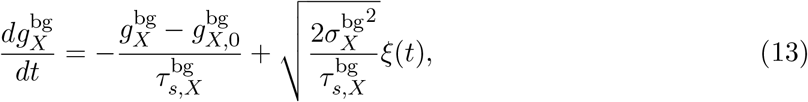

where *g*_*X*_ stands for *g*_*e*_ or *g*_*i*_, and *ξ*(*t*) is the white Gaussian noise satisfying ⟨*ξ*(*t*)⟩ = 0 and ⟨*ξ*(*t*)*ξ*(*s*)⟩ = *δ*_*ij*_*δ*(*t* − *s*).

The real biological data has a wide variety of fluctuation, including non-trivial large variations with some characteristic timescales. For instance, the hippocampal neurons are subject to the theta oscillation of the frequency range of 3 – 10 [Hz] [52]. To reproduce such oscillations that are also observed in the cross-correlogram, we introduced slow oscillations into the background noise for individual neurons, as

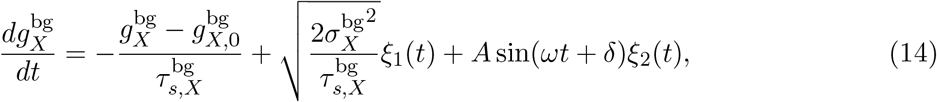

where *ξ*_1_(*t*) and *ξ*_2_(*t*) are the white Gaussian noise satisfying ⟨*ξ*_*i*_(*t*)⟩ = 0 and ⟨*ξ*_*i*_(*t*)*ξ*_*j*_(*s*)⟩ = *δ*_*ij*_*δ*(*t*−*s*).

Among *N* = 1000 neurons, we added such oscillating background signals to three subgroups of 100 neurons (80 excitatory and 20 inhibitory neurons), respectively with 7, 10, and 20 Hz. The phases of the oscillation *δ* were chosen randomly from the uniform distribution. Amplitudes of the oscillations were chosen randomly from uniform distribution in an interval [Ã*/*2, 3Ã*/*2]. The parameters for the background inputs are summarized in Table V.

**TABLE V.**
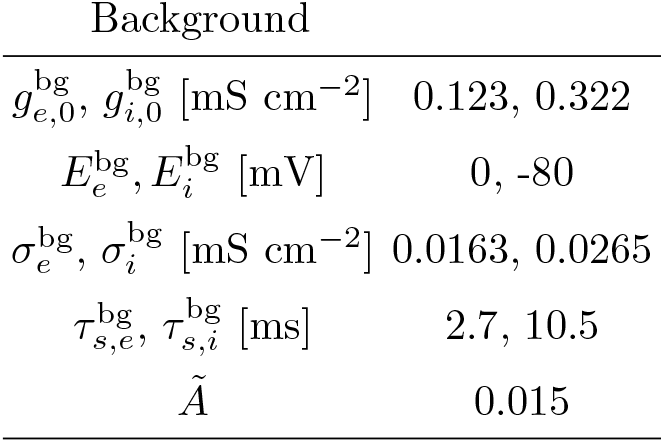
Parameters for synaptic currents and background inputs.

#### Numerical simulation

Simulation codes were written in C++ and parallelized with OpenMP framework. The time step was 0.1 ms. The neural activity was simulated up to 7,200 s.

### E. Experimental data

Spike trains were recorded from the PF, IT, and V1 cortices of monkeys in three experimental laboratories using the Utah arrays. All the studies were carried out in compliance with the ARRIVE guidelines. Individual experimental settings are summarized as follows.

#### Prefrontal cortex (PF)

The experiments were carried out on an adult male rhesus macaque *Macaca mulatta* (6.7 kg, age 4.5 y). The monkey had access to food 24 hours a day and earned liquid through task performance on testing days. Experimental monkeys were socially pair housed. All experimental procedures were performed in accordance with the ILAR Guide for the Care and Use of Laboratory Animals and were approved by the Animal Care and Use Committee of the National Institute of Mental Health (U.S.A.). Procedures adhered to applicable United States federal and local laws, including the Animal Welfare Act (1990 revision) and applicable Regulations (PL89544; USDA 1985) and Public Health Service Policy (PHS2002). Eight 96–electrode arrays (Utah arrays, 10*×*10 arrangement, 400 *µ*m pitch, 1.5mm depth, Blackrock Microsystems, Salt Lake City, U.S.A.) were implanted on the prefrontal cortex following previously described surgical procedures [53]. Briefly, a single bone flap was temporarily removed from the skull to expose the PFC, then the *dura mater* was cut open to insert the electrode arrays into the cortical parenchyma. Next, the *dura mater* was closed and the bone flap was placed back into place and attached with absorbable suture, thus protecting the brain and the implanted arrays. In parallel, a custom-designed connector holder, 3D-printed using biocompatible material, was implanted onto the posterior portion of the skull. Recordings were made using the Grapevine System (Ripple, Salt Lake City, USA). Two Neural Interface Processors (NIPs) made up the recording system, one NIP (384 channels each) was connected to the 4 multielectrode arrays of one hemisphere. Synchronizing behavioral codes from MonkeyLogic and eye-tracking signals were split and sent to each NIP. The raw extracellular signal was high-pass filtered (1kHz cutoff) and digitized (30kHz) to acquire single-unit activity. Spikes were detected online and the waveforms (snippets) were stored using the Trellis package (Grapevine). Single units were manually sorted offline using custom Matlab scripts to define time-amplitude windows in combination with clustering methods based on PCA feature extraction. Further details about the experiment can be found elsewhere [54]. Briefly, the recordings were carried out while the animals were comfortably seated in front of a computer screen, performing left or right saccadic eye movements. Each trial started with the presentation of a fixation dot on the center of the screen and the monkeys were required to fixate. After a variable time (400–800ms) had elapsed, the fixation dot was toggled off and a cue (white square, 2° *×* 2° side) was presented either to the left or right of the fixation dot. The monkeys had to make a saccade towards the cue and hold for 500ms. 70% of the correctly performed trials were rewarded stochastically with a drop of juice (daily total 175–225 mL). Typically, monkeys performed *>* 1000 correct trials in a given recording session for recording time of 120-150 minutes.

#### Inferior temporal cortex (IT)

The experiments were carried out on an adult male Japanese monkey (*Macaca fuscata*, 11 kg, age 13 y). The monkey had access to food 24 hours a day and earned its liquid during and additionally after neural recording experiments on testing days. The monkey was housed in one of adjoining individual primate cages that allowed social interaction. All experimental procedures were approved by the Animal Care and Use Committee of the National Institute of Advanced Industrial Science and Technology (Japan) and were implemented in accordance with the “Guide for the Care and Use of Laboratory Animals” (eighth ed., National Research Council of the National Academies). Four 96 microelectrode arrays (Utah arrays, 10 *×* 10 layout, 400 *µ*m pitch, 1.5 mm depth, Blackrock Microsystems, Salt Lake City, USA) were surgically implanted on the IT cortex of the left hemisphere. Three arrays were located in area TE and the remaining one in area TEO. Surgical procedures were roughly the same as having been described previously [53], except that a bone flap that was temporarily removed from the skull was located over the IT cortex and that a CILUX chamber was implanted onto the anterior part of the skull protecting connectors of the arrays. Recordings of neural data and eye positions were done in a single session using Cerebus™ system (Blackrock Microsystems). The extracellular signal was band-pass filtered (250–7.5 k Hz) and digitized (30 kHz). Units were sorted online before the recording session for the extracellular signal of each electrode using a threshold and time-amplitude windows. Both the spike times and the waveforms (10 and 38 samples, preceding and after a threshold crossing, respectively) of the units were stored using Cerebus Central Suite (Blackrock Microsystems). Single units were refined offline by hand using the PCA projection of the spike waveforms in Offline sorter™ (Plexon Inc., Dallas, USA). The monkey seated in a primate chair, and the head was restrained with a head holding device so that the eyes were positioned 57 cm in front of a color monitor’s display (GDM-F520, SONY, Japan). The display subtended a visual angle of 40° *×* 30° with a resolution of 800 *×* 600 pixels. A television series on animals (NHK’s Darwin’s Amazing Animals, Asahi Shimbun Publications Inc., Japan) was shown on the display throughout the online spike sorting and the recording session. The monkey’s eye position was monitored using an infrared pupil-position monitoring system [55] and was not restricted.

#### The primary visual cortex (V1)

The data set was obtained from Collaborative Research in Computational Neuroscience (CR-CNS), pvc-11 [56] by the courtesy of the authors of [57]. In this experiment, spontaneous activity was measured from the primary visual cortex while a monkey viewed a CRT monitor (1024 *×* 768 pixels, 100 Hz refresh) displaying a uniform gray screen (luminance of roughly 40 cd/m^2^). Briefly, the animal was premedicated with atropine sulfate (0.05 mg/kg) and diazepam (Valium, 1.5 mg/kg) 30 min before inducing anesthesia with ketamine HCl (10.0 mg/kg). Anesthesia was maintained throughout the experiment by a continuous intravenous infusion of sufentanil citrate. To minimize eye movements, the animal was paralyzed with a continuous intravenous infusion of vecuronium bromide (0.1 mg/kg/h). Vital signs (EEG, ECG, blood pressure, end-tidal PCO2, temperature, and lung pressure) were monitored continuously. The pupils were dilated with topical atropine and the corneas protected with gas-permeable hard contact lenses. Supplementary lenses were used to bring the retinal image into focus by direct ophthalmoscopy and later adjusted the refraction further to optimize the response of recorded units. Experiments typically lasted 4–5 d. All experimental procedures complied with guidelines approved by the Albert Einstein College of Medicine of Yeshiva University and New York University Animal Welfare Committees.

Spike sorting and analysis criteria: Waveform segments were sorted off-line with an automated sorting algorithm, which clustered similarly shaped waveforms using a competitive mixture decomposition method [58]. The output of this algorithm was refined by hand with custom time-amplitude window discrimination software (written in MATLAB; MathWorks) for each electrode, taking into account the waveform shape and interspike interval distribution. To quantify the quality of the recording, the signal-to-noise ratio (SNR) of each candidate unit was computed as the ratio of the average waveform amplitude to the SD of the waveform noise [59–61]. Candidates that fell below an SNR of 2.75 were discarded as multiunit recordings.

## ACKNOWLEDGMENTS

We thank Masahiro Naito for his technical assistance in developing a web-application program, Jun-nosuke Teramae for his advice on MAT simulation, and Kai Shinomoto for drawing an illustration of a monkey for Figure 1. We also thank Adam Kohn for permitting us to analyze their experimental data of V1 and providing the detailed information of the experimental conditions, and Richard Saunders and Mark Eldridge for performing surgery on the animal for IT cortex data, Yuji Nagai and Takafumi Minamimoto for assisting the surgery, and Rossella Falcone and Narihisa Matsumoto for helpful discussions upon preparing the IT cortex data.

R.K. is supported by JSPS KAKENHI Grant Numbers JP17H03279, JP18K11560, JP19H01133, and JPJSBP120202201, and JST PRESTO Grant Number JPMJPR1925, Japan. B.B.A. is supported by NIMH DIRP ZIA MH002928. Y.S.M is supported by JSPS KAKENHI Grant Number JP18H05020 and New Energy and Industrial Technology Development Organization (NEDO). K.H. is supported by Japan Society for the Promotion of Science (JSPS) and JSPS KAKENHI Grant Number JP19J40302. K.K. is supported by JSPS KAKENHI Grant Number JP19K07804. B.J.R. is supported by NIMH DIRP ZIA MH002032. S.S. is supported by JST CREST Grant Number JPMJCR1304, and the New Energy and Industrial Technology Development Organization (NEDO).

